# Analysing plasma membrane asymmetry of lipid organisation by fluorescence lifetime and correlation spectroscopy

**DOI:** 10.1101/776831

**Authors:** Anjali Gupta, Thomas Korte, Andreas Herrmann, Thorsten Wohland

## Abstract

A fundamental feature of a eukaryotic cell membrane is the asymmetric arrangement of lipids in the two leaflets. A cell invests significant energy to maintain this asymmetry and utilizes it to regulate important biological processes such as apoptosis and vesiculation. Here, we employ *fluorescence lifetime imaging microscopy* (FLIM) and *imaging total internal reflection fluorescence correlation spectroscopy* (ITIR-FCS) to differentiate the dynamics and organization of the exofacial and cytoplasmic leaflet of live mammalian cells. We characterize the biophysical properties of fluorescent analogues of phosphatidylcholine (PC), sphingomyelin (SM) and phosphatidylserine (PS) in two mammalian cell membranes. Due to their specific transverse membrane distribution, these probes allow leaflet specific investigation of the plasma membrane. We compare the results with regard to the different temporal and spatial resolution of the methods. Fluorescence lifetimes of fluorescent lipid analogues were found to be in a characteristic range for the liquid ordered phase in the outer leaflet and liquid disordered phase in the inner leaflet. The observation of a more fluid inner leaflet is supported by free diffusion in the inner leaflet with high average diffusion coefficients. The liquid ordered phase in the outer leaflet is accompanied by slower diffusion and diffusion with intermittent transient trapping. Our results show that the combination of FLIM and ITIR-FCS with specific fluorescent lipid analogues provides a powerful tool to investigate lateral and trans-bilayer characteristics of plasma membrane in live cells.

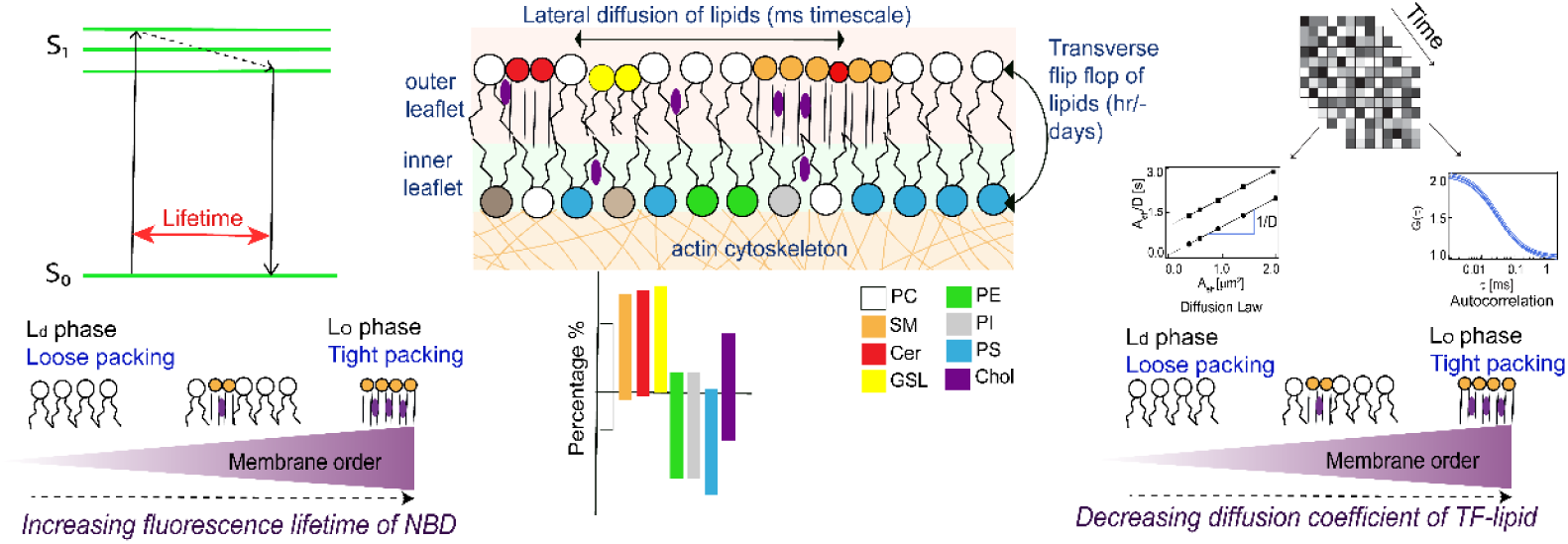

## INTRODUCTION

The plasma membrane is a bilayer composed of a plethora of chemically diverse lipids and a range of proteins. The distinct physicochemical properties of the membrane components lead to the formation of transient assemblies such as cholesterol-dependent domains, cholesterol independent domains and protein oligomers which are typically below the diffraction limit (20-100 nm) and are in dynamic equilibrium with each other^1,2,3^. The transient assemblies exhibit unique lifespans governed by the mutual interactions of their components. In addition to the lateral heterogeneity, a fundamental feature of the plasma membrane is the asymmetry of lipid composition between the two leaflets^4–6^. The dynamic lateral heterogeneity of the membrane is essential for cell signalling^7^ while the asymmetric arrangement of lipids across the plasma membrane is found to be critical in the regulation of several biological processes, including apoptosis^8^, cell-cell fusion^9^ and signalling in immune cells^10^. The primary classes of lipids that constitute a typical mammalian plasma membrane are glycerophospholipids (e.g. PC, PS, PE, PA, PI), sphingolipids (e.g. SM, ceramide, GSLs) and cholesterol^11^. In the plasma membrane, PC and sphingolipids are prevalent in the outer leaflet while aminophospholipids such as PS, and PE are prevalent in the inner leaflet^4^. Since the outer-leaflet comprises more domain forming lipids and the inner-leaflet is in direct contact with the cytoskeleton, it is expected that the organization and dynamics of the two leaflets are different. However, the precise lateral and transbilayer organization of the membrane and how the two leaflets are dynamically coupled are unknown. Several studies have focused on understanding membrane asymmetry of organisation, however, due to experimental limitations they have mostly been conducted on model membranes with limited physiological relevance as it is difficult to recapitulate the structural complexity of an intact cell membrane^12–14^. A detailed analysis of plasma membrane asymmetry in physiological context requires a methodology that can probe dynamics and organization of both leaflets of an intact cell membrane separately with sufficient spatiotemporal resolution. In addition to the quantitative nature of the method, the other key points necessary to analyze the transbilayer membrane organization are^15^: (1) There should be a way of separating the information originating from the two leaflets. This can be achieved by using lipid probes that confine themselves predominantly to one particular leaflet or by selectively quenching the fluorescence signal from one leaflet using a membrane impermeable agent, e.g. sodium dithionite, TNBS, and Doxyl-PC. (2) The time taken to complete the assay should be less than what it takes for lipids to undergo transbilayer movement. (3) The quenching agent should not have any additional effects on the membrane integrity. (4) Certain chemical treatments can alter the membrane lipid composition by modulating the rates of endo-or exocytosis. During the analysis of plasma membrane asymmetry, any chemical treatment should not alter the membrane composition.

Based on the above-mentioned considerations, we perform a detailed analysis of the plasma membrane asymmetry by examining the outer and the inner leaflet of the membrane separately in live mammalian cells using a combination of *fluorescence lifetime imaging microscopy (FLIM)* and *imaging total internal reflection fluorescence correlation spectroscopy (ITIR-FCS)*. FLIM has been utilized to investigate the asymmetrical arrangement of the plasma membrane^16,17^. It can reliably detect and resolve microscopic domains present in model membranes as identified by their different lifetimes. However, in GPMVs and intact plasma membranes, where lateral heterogeneities are on the nanometer scale, FLIM is unable to resolve discrete membrane domains. Due to the domains, FLIM detects a mixture of lifetimes leading to a broad distribution of fluorescence lifetimes indicating the existence of a wide variety of transient assemblies^16^. In order to obtain complementary information to membrane organization as measured by FLIM we use ITIR-FCS, to measure membrane dynamics^18–21^. ITIR-FCS is an imaging modality that allows multiplexed FCS measurements on the whole region of interest simultaneously and provides spatially resolved diffusion coefficient and number of particle maps. And although ITIR-FCS is diffraction-limited, the use of the FCS diffusion law, which determines the change of diffusion with spatial scale, allows the investigation of sub-resolution membrane organization ^22,23^. FLIM measurements require environment-sensitive fluorophores while for FCS measurements fluorophores with better photostability are ideal. Therefore, we use 1-palmitoyl-2-[6-[(7-nitro-2-1,3-benzoxadiazol-4-yl)amino]-hexanoyl]-sn-glycero-3-phospholipid (NBD) labeled lipid analogues for FLIM as NBD lifetime is sensitive to the environmental polarity and molecular packing around the probe^17,24–26^ (Figure 1*A*). However, low photostability and brightness of NBD make it non-ideal for FCS measurements^27^. Therefore, we use dipyrrometheneboron difluoride (TopFluor) labeled lipids, which offer sufficient photostability, for FCS measurements (Figure 1*A*). To evaluate cell type, domain, and leaflet specific information, we examine the biophysical properties of fluorescently labeled PC, SM, and PS analogues in the plasma membrane of CHO-K1 and RBL-2H3 cells. Our FLIM results show that the lifetime of all the tested probes is typically longer in the outer leaflet than in the inner leaflet most convincingly shown for NBD-PS which is predominantly localized in the inner leaflet of the membrane. In support of lifetime results, ITIR-FCS measurements on the same cell lines show a higher diffusion coefficient and less domain confined diffusion of lipid probes localized in the inner leaflet of the membrane and a slower diffusion accompanied with higher confinement in the outer leaflet of the membrane, with characteristic differences between cell lines. Overall, our results reveal that the outer leaflet exhibits a liquid-ordered environment while the inner leaflet of the membrane is more similar to the liquid disordered phase. Furthermore, we show cell line differences in the transbilayer organization of the plasma membrane.

**Figure 1:**
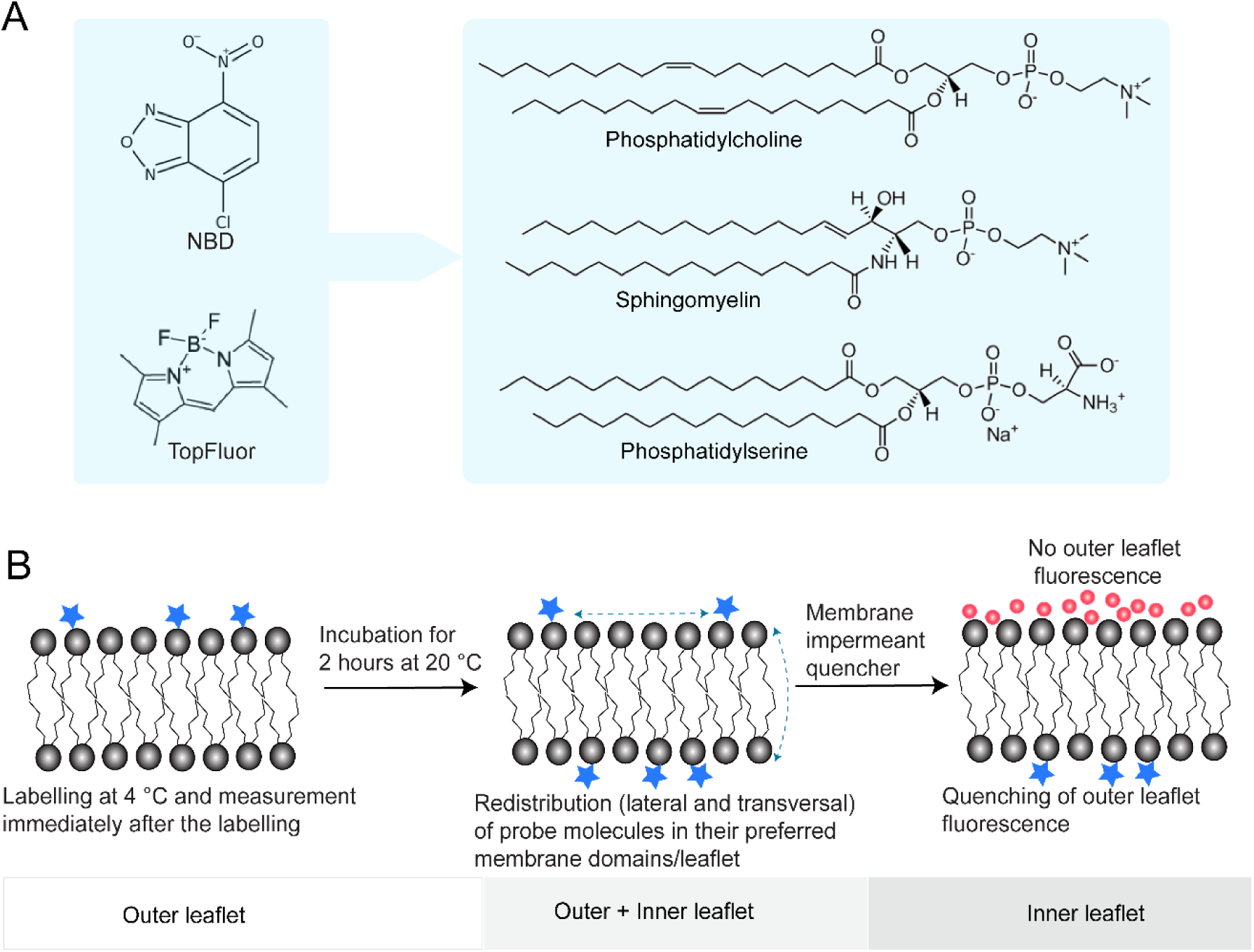
(A) Chemical structures of the lipid analogs used in this study. 1-palmitoyl-2-[6-[(7-nitro-2-1,3-benzoxadiazol-4-yl) amino]-hexanoyl]-sn-glycero-3-phospholipid (NBD) and dipyrrometheneboron difluoride (TopFluor) labeled PC, SM and PS analogues. (B) For the readouts from outer leaflet exclusively, cells are labeled at 4 °C and are measured immediately. Then probes (lipids marked with asterisks) are allowed to redistribute in the cell membrane via both lateral and transversal movement, and fluorescence is recorded from probes localized in both outer and inner leaflets. Finally, outer leaflet probe fluorescence is quenched by treating the labeled cells with membrane impermeant quencher as shown by red circles (NBD: sodium dithionite; TopFluor: Doxyl-16 lipids) at 4 °C, and then the measurements are performed. Post quenching readouts provide access to inner leaflet organization and dynamics exclusively.

## EXPERIMENTAL SECTION

### Material and Sample preparation

1-palmitoyl-2-{6-[(7-nitro-2-1,3-benzoxadiazol-4-yl)amino]hexanoyl}-sn-glycero-3-phosphocholine (NBD-PC), N-[6-[(7-nitro-2-1,3-benzoxadiazol-4-yl)amino]hexanoyl]-sphingosine-1-phosphocholine (NBD-SM), 1-palmitoyl-2-{6-[(7-nitro-2-1,3-benzoxadiazol-4-yl)amino]hexanoyl}-sn-glycero-3-phosphoserine (ammonium salt) (NBD-PS), 1-palmitoyl-2-(dipyrrometheneboron difluoride) undecanoyl-sn-glycero-3-phosphocholine (TF-PC), N-[11-(dipyrrometheneboron difluoride)undecanoyl]-D-erythro-sphingosylphosphorylcholine (TF-SM) and 1-palmitoyl-2-(dipyrrometheneboron difluoride) undecanoyl-sn-glycero-3-phospho-L-serine (ammonium salt) (TF-PS) (Figure 1A), 1-palmitoyl-2-stearoyl-(16-doxyl)-sn-glycero-3-phosphocholine (16:0-16 Doxyl PC) were purchased from Avanti Polar Lipids (Alabaster, AL). Lipid analogs were solubilized in methanol. Sodium dithionite was purchased from Sigma-Aldrich, Singapore. Chinese hamster ovary (CHO-K1) and rat basophilic leukemia (RBL-2H3) cells – were purchased from ATCC (Manassas, VA). We use NBD lipid analogs for discerning the microenvironments of the conjugated lipids in CHO-K1 and RBL-2H3 cell membranes by performing FLIM experiments. The fluorescence lifetime of NBD is influenced by the surrounding membrane packing, environmental polarity and tendency to show red edge excitation shift (REES). Here, we use TopFluor lipid conjugates for ITIR-FCS measurements as previous reports demonstrate their compatibility with FCS method^28^. Sodium dithionite and Doxyl PC are used to quench outer leaflet NBD and TopFluor lipid analogues respectively. Prior to FLIM measurements on cell membranes, lifetime measurements were performed on large unilamellar vesicles (LUV) labeled with NBD-PC at temperature range (25 °C - 45 °C).

### Large unilamellar vesicle preparation

LUVs were prepared by evaporating the chloroform from the calculated amounts of lipids and the fluorescent lipid analogue in a glass flask under a stream of nitrogen. Dried lipid was resuspended in the buffer solution (150 mM NaCl, 10 mM HEPES, pH 7.4). Then the solution was subjected to freeze-thaw cycle for at least eight times to obtain multilamellar vesicles. Finally, the multilamellar vesicle solution was extruded at least ten times through 100 nm polycarbonate filters (Whatman Schleicher & Schnell) at a temperature above lipid phase separation. For fluorescence lifetime measurements we used mM total lipid LUV-solutions.

### Giant plasma membrane vesicle preparation

Giant plasma membrane vesicles (GPMVs) were prepared from CHO-K1 and RBL-2H3 cells by the formaldehyde-based induction of plasma membrane blebbing. The detailed protocol is described in Sezgin et al^40^.

### Cell membrane labeling

The stock of both NBD and TopFluor lipid analogues (PC, SM and PS) (Avanti Polar Lipids (Alabaster, AL) was prepared in pure methanol. For cell membrane staining with NBD lipid analogues, stock solution was diluted in HBSS (Hank’s Balanced Salt Solution; Invitrogen, Singapore) to a final concentration of 4 μM. The diluted solution was added to the cells followed by an incubation for 20 minutes on ice. After incubation, the cells were washed with HBSS at least three times and then cells were used for FLIM experiments. For cell membrane staining with TopFluor lipid analogues, stock solution was vigorously vortexed and was diluted in HBSS to a final concentration of 5 μM for CHO-K1 cells and 2 μM for RBL-2H3 cells. 3mg/mL bovine serum albumin (Sigma Aldrich, Singapore) was added to the working solution of the fluorescent lipid analogues. The cells were incubated with the working solution (lipid analogue:BSA) at 4 °C for 15-20 minutes. After incubation, the cells were washed with imaging medium (DMEM without phenol red (Invitrogen, Singapore) and 10 % FBS) at least three times and then the cells were used for imaging.

### Preparation of quenching solutions for NBD and TopFluor conjugated lipid probes

The outer leaflet fluorescence of NBD and TopFluor was quenched by sodium dithionite and 16-Doxyl-PC respectively. A stock solution of 1M sodium dithionite was prepared. Then the stock solution was diluted to a concentration of 25mM in 10mM Tris (pH 9) and was used as the working solution.16-Doxyl-PC was solubilised in pure methanol. Then calculated amount of 16-Doxyl-PC was added to HBSS to a final concentration of 16 μM.

### Determination of the percentage of fluorescent lipid analogues present in the outer leaflet using fluorescence spectroscopy

The fraction of fluorescent lipid analogues was evaluated by the quenching of outer leaflet fluorescence selectively using cell impermeant fluorescence quenchers. Cell suspension of about 10^6^ cells was centrifuged at 1000 rpm for 5 minutes. Then pelleted cells were suspended in the working solution of fluorescent lipid analogues and were incubated for 15 minutes. Cells were again pelleted by centrifugation and were resuspended in HBSS. For the redistribution of lipid probes, cells were incubated at room temperature for 2 hours. The fluorescence intensity of these cells was measured (excitation at 470 nm) on Cary Eclipse Fluorescence Spectrophotometer, Singapore. Then cells were pelleted down and HBSS was replaced with the quenching solution. Cells were suspended in cold quenching solution and were incubated on ice for 10 minutes. Cold conditions prevent the permeability of quencher inside the cell. Then cells are washed with HBSS at least thrice and were used for fluorescence intensity measurements.

### Cholesterol depletion experiment

Cholesterol depletion was performed by treating the cells with 3 mM methyl-β-cyclodextrin (Sigma Aldrich, Germany). The cells were incubated for 30 minutes were then measured.

### Fluorescence lifetime spectroscopy

Fluorescence lifetime spectroscopy (FLS) was performed on a FluoTime 200 spectrometer (PicoQuant, Germany). A pulsed laser diode (LDH-P-C, PicoQuant, Germany) of 470 nm and a pulse frequency of 8 MHz was used as the excitation source for NBD probes. Individual photons were recorded on a time correlated single photon counting (TCSPC) setup with a time resolution of 33 ps. NBD emission was recorded at 540 nm and the spectral bandwidth was set to 4 nm. Mean photon count rates were kept ∽ 1-4 × 10^4^ counts/s. The instrument response function (IRF) was recorded using HEPES buffered saline (HBS) at excitation wavelength. Intensity decays were fitted globally (over at least 7 data sets) using a nonlinear least square iterative reconvolution fitting procedure (FluoFit, PicoQuant, Germany):

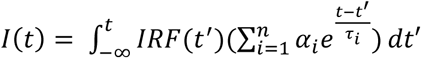

where I(t) is the fluorescence intensity at time t, α_i_ the pre-exponential factor representing the intensity of the time-resolved decay of the component with lifetime τ_i_. All intensity decays were fitted to bi- or triexponential model functions depending on the studied sample. We ensured the quality of fit by the χ^2^ value, the distribution of residuals and the autocorrelation function of residuals. The fitting error was calculated using a support plane error analysis and included into the error estimation.

#### Fluorescence lifetime imaging microscopy

##### Instrumentation and data acquisition

Images were acquired by a confocal laser scanning microscope with an inverted Fluoview 1000 microscope (Olympus, Tokyo, Japan) and a 60X (NA 1.35) oil-immersion objective at 25 °C. The frame size of the acquired images is 512 × 512 pixels. For images of cells labeled with NBD fluorescent analogues, the fluorophores were excited with a 488 nm argon ion laser and the signal was recorded between 500 and 530 nm. For the recording of FLIM measurements we used a commercial FLIM upgrade kit (PicoQuant, Berlin, Germany). For these measurements, NBD was excited with a pulsed diode laser (pulse width: 60 ps; pulse frequency: 10 MHz; 4 μs/pixel) with a wavelength of 483 nm. Emission light was filtered using a 540/40 bandpass filter and then single photons were registered with a single photon avalanche photo diode.

For each FLIM measurement 50–70 frames were recorded; the average photon count rate was kept around ∼2–4 × 10^4^ counts/s. The images were pseudocolor-coded in accordance with the average lifetime (τ_av_) of the pixels.

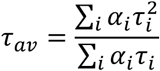

##### Data analysis

For extracting the fluorescence lifetimes of NBD analogues, membrane regions are selected by applying an intensity threshold to exclude fluorescence from background or cytoplasm. The selection was further refined manually to exclude regions not associated with the membrane. Then overall fluorescence decay curve was calculated by adding up the photons registered from the selected region. The part of the decay curve that comprises instrument response function was removed, and the rest was used for the analysis. A non-linear least squares iterative fitting procedure was used to fit the fluorescence decay curves as a sum of exponential terms,

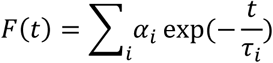

where F(t) denotes the fluorescence intensity at time t, and *α*_*i*_ denotes a pre-exponential factor representing the intensity of the time-resolved decay of the component with lifetime *τ*_*i.*_ The quality of fits was evaluated by the distribution of the residuals and the χ^2^ value.

A typical FLIM experiment yields a spatial distribution of lifetime (usually lifetime is mapped in a color-coded fashion) however since the size of membrane domains is below the spatial resolution individual images do not provide any additional insights. Thus, we represent the data in the form of lifetimes and amplitude averaged over the whole membrane.

#### Imaging total internal reflection fluorescence correlation spectroscopy

The experiments were done on TopFluor lipid analogue labeled cell membranes. The experiments are performed at 25 °C with 5% CO_2_.

(a) Imaging Total internal reflection fluorescence correlation spectroscopy (ITIR-FCS)

ITIR-FCS measurements were conducted on an objective type TIRF microscope (IX-71, Olympus, Singapore). The current study uses a high NA oil immersion objective (PlanApo, 100×, NA 1.45, Olympus, Singapore) and the sample is excited using a 488 nm laser (Spectra-Physics Lasers, Mountain View, CA, US) which was then directed into the microscope by a combination of two tilting mirrors. The laser power used for all the experiments lies in the range of 0.8-1mW. Then the light is reflected by a dichroic mirror (Z488/532RPC, Semrock) and was focused on the objective back focal plane. The incident angle of the light is controlled by the same combination of tilting mirror and it was total internally reflected at the glass-water interface. Finally, the light is filtered by the emission filter and was detected on the CCD chip of a cooled (−80 °C), back-illuminated EMCCD camera (Andor iXON 860, 128×128 pixels, Andor technology, US). Data acquisition is performed using Andor Solis software (version 4.18.30004.0 and 4.24.30004.0). The pixel side length of the CCD chip in the device is 24 μm corresponding to a pixel side length of 240 nm in the sample plane. During data acquired in the kinetic mode, and to reduce the baseline fluctuations baseline clamp was switched on. The readout speed was 10 MHz with 4.7x maximum analog-to-digital gain and 25 μs vertical shift speed. An EM gain of 300 was used for all the ITIR-FCS experiments.

The fluorescence intensity signal was recorded from a 21 × 21 pixels region of interest (ROI) simultaneously as a stack of 30,000 - 50,000 frames with 2 ms time resolution. The data was saved as 16-bit Tiff file. The fluctuations contained in the temporal intensity trace from each pixel was autocorrelated using multi-tau correlation scheme using a FIJI plug-in ImFCS 1.49, a home-written software which is provided at this link xshttp://www.dbs.nus.edu.sg/lab/BFL/imfcs_image_j_plugin.html] to obtain autocorrelation functions (ACF)^58^. Bleach correction is applied on the data using a 4^th^ order polynomial function. The ACF for each pixel was individually fitted with the following one-particle model for diffusion using the same software.

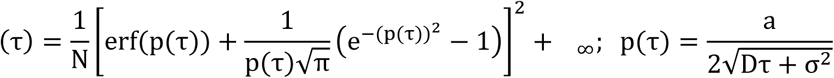

Here *G(τ)* represents the ACF as a function of correlation time (*τ*) and *N, a, D* and *σ* are the number of particles per pixel, pixel side length, diffusion coefficient and standard deviation of the Gaussian approximation of the microscope point spread function (PSF) respectively. *G*_∞_ represents the convergence value of the ACF at long correlation times.

Fitting of experimentally obtained ACFs with theoretical models yields *D* and *N*. Since it is an imaging based FCS modality, we obtain spatially resolved diffusion coefficient (*D*), and the number of particles (*N*) maps^58^. In this study, the data are represented as mean ± standard deviation (SD). The SD is obtained from the measurements over 441 pixels per experiment. The SD of an ITIR-FCS measurement not only contains contributions from the measurement variability but also from the cell membrane heterogeneity. Each data point for diffusion coefficient obtained using ITIR-FCS is an average of 441 measurements therefore, N = 5292 at least.

##### Imaging FCS diffusion laws

For the analysis of cell membrane organization below the diffraction limit, we combined ITIR-FCS with FCS diffusion laws. These laws allow probing mode of molecular diffusion i.e. free diffusion or is hindered by the trapping sites such as domains^22,23^. To achieve this, we plot the spatial dependence of the diffusion time of the probe molecules on the observation area size. Then these plots are fitted to a standard error of mean (SEM) weighted straight line which is mathematically expressed as:

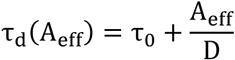

Where τ_0_ represents the FCS diffusion law intercept. For deriving sub-resolution information, the diffusion law plot is extrapolated to zero and y-intercept (*τ*_0_) is used as an indicator of membrane organization. For a freely diffusing particle, the diffusion time scales linearly with the observation area and the y-intercept is zero. In case of domain entrapment and hop diffusion, the relationship between the diffusion time of the molecules and the observation area becomes non-linear and the obtained y-intercept is positive and negative respectively. To perform FCS Diffusion Law analysis, the data that is acquired in an ITIR-FCS experiment is utilized. Post-acquisition pixel binning (1×1 to 5×5) followed by convolution with the PSF of the microscope system to generate variable observation areas (*A*_eff_). The typical margin of error on cell membrane is ±0.1 and thus intercepts in that lie in this range are indistinguishable for free diffusion. Only intercepts greater than 0.1 can be attributed to domain entrapment in our setup.

## RESULTS AND DISCUSSIONS

Here, we investigated the dynamics and organization of the outer and the inner leaflet in the plasma membrane of mammalian cells. For outer leaflet membrane dynamics, we performed measurements at room temperature immediately after labeling of cells with fluorescent lipid analogues at 4 °C. Both FLIM and ITIR-FCS measurements are done at room temperature within the first 1-5 minutes after staining. As this is less than the time needed for the transbilayer movement of glycerophospholipids, which typically ranges from several minutes to hours at 37°C ^29^, measured values report essentially on the outer leaflet. Subsequently, we measure after an incubation of 2 hours at room temperature. During this time, lipid analogues redistribute to their preferred locations in the membrane^30,31^. Aminophospholipids (eg, PS) are recognized by ATP dependent flippases and are rapidly translocated to the inner leaflet of the membrane^32^. In addition, smaller percentages of choline lipids are also translocated to the inner leaflet via passive diffusion. At this stage, we obtain a combined readout of both outer and inner leaflet. Finally, for the exclusive measurement of inner leaflet dynamics and organization, outer leaflet fluorescence is quenched. The selective quenching of outer leaflet fluorescence is achieved by treating labeled cells by a suitable membrane impermeable quencher at 4 °C, (Figure 1*B*). NBD lipid analogues are reduced to the non-fluorescent product 7-amino-2,1,3-benzoxadiazol-4-yl (ABD) by treating the labeled cells with sodium dithionite and TopFluor lipid analogues are quenched by spin-labeled 16-Doxyl PC^28,30,31^. The quenching efficiency has been quantified for each fluorescent lipid analogue in both CHO-K1 and RBL-2H3 cell membranes (Table S1).

FLIM measures the fluorescence lifetimes and their fraction, which report on the packing in the probe microenvironment and the prevalence of molecules in different environments, respectively. In an ITIR-FCS experiment, membrane dynamics is characterized by the diffusion coefficient (*D*) and membrane organization by the FCS diffusion law intercept (*τ*_*0*_). We use the diffusion coefficient as a readout of molecular mobility of the probe molecules residing in the membrane, and it is likely to increase with a decrease in membrane domain fraction. The FCS diffusion law intercept (see Material and Method section) provides information about the mode of probe organization in the membrane estimated by plotting the dependence of diffusion time over observation area size^22,23^. For a freely diffusing particle, the diffusion law intercept lies in the range of -0.1 s < τ_0_ < 0.1s. A positive intercept (τ_0_ > 0.1s) represents transient entrapment of the probe, e.g., in cholesterol sphingomyelin complexes, and a negative intercept (τ_0_ < -0.1s) implies a meshwork organization and hop diffusion of probe molecules, e.g., molecules hindered by the cytoskeleton. In situations where a molecule is transiently trapped and hindered by a meshwork simultaneously, the intercept represents a weighted average of both diffusion modes^33^. In this case, additional measurements that influence either one of the diffusion modes are necessary to disentangle the contributions by hop diffusion and transient trapping.

### The fluorescence lifetime of NBD-PC in membranes with discrete domains

First we determined the NBD lifetime within different membrane phases, by measuring the lifetime of NBD-PC in large unilamellar vesicles (LUVs) with defined lipid compositions and known phase behaviours^34,35^. The measurements are performed at a temperature range of 25 - 45 °C. DOPC/SSM/Chol LUVs with molar ratios of 2:2:6, and 1:1:1 form a pure liquid ordered (l_o_) phase or coexisting l_d_ - l_o_ phases, respectively. For a pure liquid disordered (l_d_) phase, DOPC LUVs are used. The fluorescence decay of NBD in LUVs exhibiting a single phase is best fitted to a bi-exponential model (Figure S1). Past studies have suggested that only the long lifetime is sensitive to the membrane environment while the short lifetime is not dependent on the membrane phase^16,24^. Lifetime data from LUVs with coexisting phases is best fitted to a tri-exponential model where the two long lifetimes represent the two membrane phases (l_d_ and l_o_), and the third, the short component is not sensitive to membrane phases. The information about the lifetime of NBD in discrete domains with known phases provides an expected range of lifetime in the membrane environment and aids in better interpretation of cell membrane data. Since our cell membrane experiments are performed at 25 °C it is important to take a closer look at the lifetimes originating from the two extreme phases i.e. l_o_ and l_d_ phase at this temperature. At 25 °C, NBD lifetime in the l_d_ phase, i.e. in pure DOPC LUVs, is 6.80 ± 0.04 ns and in the l_o_ phase, i.e. in DOPC:SSM:Chol (2:2:6), is 9.94 ± 0.05 ns. LUVs with coexisting domains i.e. DOPC:SSM:Chol (1:1:1), show two membrane environment sensitive lifetimes, 9.62 ± 0.2 ns in l_o_ phase and 5.06 ± 0.3 ns in l_d_ phase. For a detailed description of the obtained results refer to *Supplementary Information S1*.

In the next sections, we present the domain and leaflet specific dynamics and organization of fluorescent lipid analogues in CHO-K1 and RBL-2H3 cell membranes, which are known to manifest distinct membrane properties and composition^36,37^.

### Leaflet specific analysis of phosphatidylcholine fluorescent analogues in CHO-K1 and RBL-2H3 cell membranes

Phosphatidylcholine (PC) is the most abundant lipid in most mammalian membranes. The molecular geometry of PC is cylindrical, and it has the capability of self-organizing as a bilayer. Most PC molecules present in cell membranes are fluid at room temperature^11^. More than 50% of the PC lipid analogues reside in the outer leaflet of the plasma membrane as observed by the estimated quenching efficiency (Table S1). The analysis of NBD-PC fluorescence lifetimes in the outer leaflet of CHO-K1 and RBL-2H3 cells reveal a lifetime of about 10 ns in both cell lines (Figure 2 *A, B*) along with the percentage fraction of long lifetime component ranging between 40-55%. By comparing the NBD-PC lifetime obtained in cell membranes with those in LUVs, it is evident that the outer leaflet of both plasma membranes contains a fraction of l_o_ phase domains. Using the same experimental conditions, ITIR-FCS measurements on TF-PC show a two-fold faster *D* (Figure 2 *C, D*) in CHO-K1 cells (*D* = 0.55 ± 0.15 μm^2^/s) as compared to RBL-2H3 cells (*D* = 0.24 ± 0.03 μm^2^/s). In addition, it also reveals a transient entrapment as indicated by a positive *τ*_*0*_ for this probe in both cell lines with RBL-2H3 cells (*τ*_*0*_ =1.48 ± 0.23 s) showing a higher value than CHO-K1 cells (*τ*_*0*_ = 0.78 ± 0.27 s). Subsequently, labeled cells were measured again after the redistribution of probes between both membrane leaflets (see Fig. 1B, middle). These results contain information averaged over both leaflets. Due to technical limitations, the leaflet specific lipid composition is difficult to analyze in cell membranes. However, it has been reported that the asymmetric distribution of PC differs across cell lines^38^. FLIM analysis of NBD-PC labeled cells post redistribution reveals no significant changes in the probe lifetime as it remains around 10 ns (t-test; p value > 0.05). However, ITIR-FCS demonstrates a 30-40 % higher *D* in both cell lines and a lower *τ*_*0*_ (CHO-K1: 0.78 ± 0.27 s to 0.64 ± 0.41 s; RBL-2H3: 1.48 ± 0.23 s to 1.14 ± 0.17 s) indicating differences in the inner leaflet organization of both the cell types. Thus, for a clearer understanding of the inner leaflet organization we measure the biophysical properties of this probe exclusively on the inner leaflet of both cell lines by quenching fluorescent lipids in the outer leaflet. In this case, FLIM reveals a 2 ns reduction in the longest lifetime component as compared to that measured on the outer leaflet. With reference to the LUV data (Figure S1), this suggests that inner leaflet resembles an l_d_ phase. This is supported by the ITIR-FCS results that show a lower *D* in the outer leaflet relative to inner leaflet (CHO-K1 : 0.55 μm^2^/s to 1.11 μm^2^/s; RBL-2H3 : 0.24 ± 0.03 μm^2^/s to 0.43 ± 0.25 μm^2^/s) and concurrent drop in *τ*_*0*_ (CHO-K1 : 0.78 ± 0.27 s to 0.25 ± 0.06 s; RBL-2H3 : 1.48 ± 0.23 s to 0.85 ± 0.22 s). As reflected by a higher *D* and lower *τ*_*0*_ in both cell lines, these results indicate that the liquid ordered domain fraction is lower in the inner leaflet. When comparing the two cell lines, the domain fraction is more elevated in RBL-2H3 than CHO-K1 inner leaflets.

**Figure 2:**
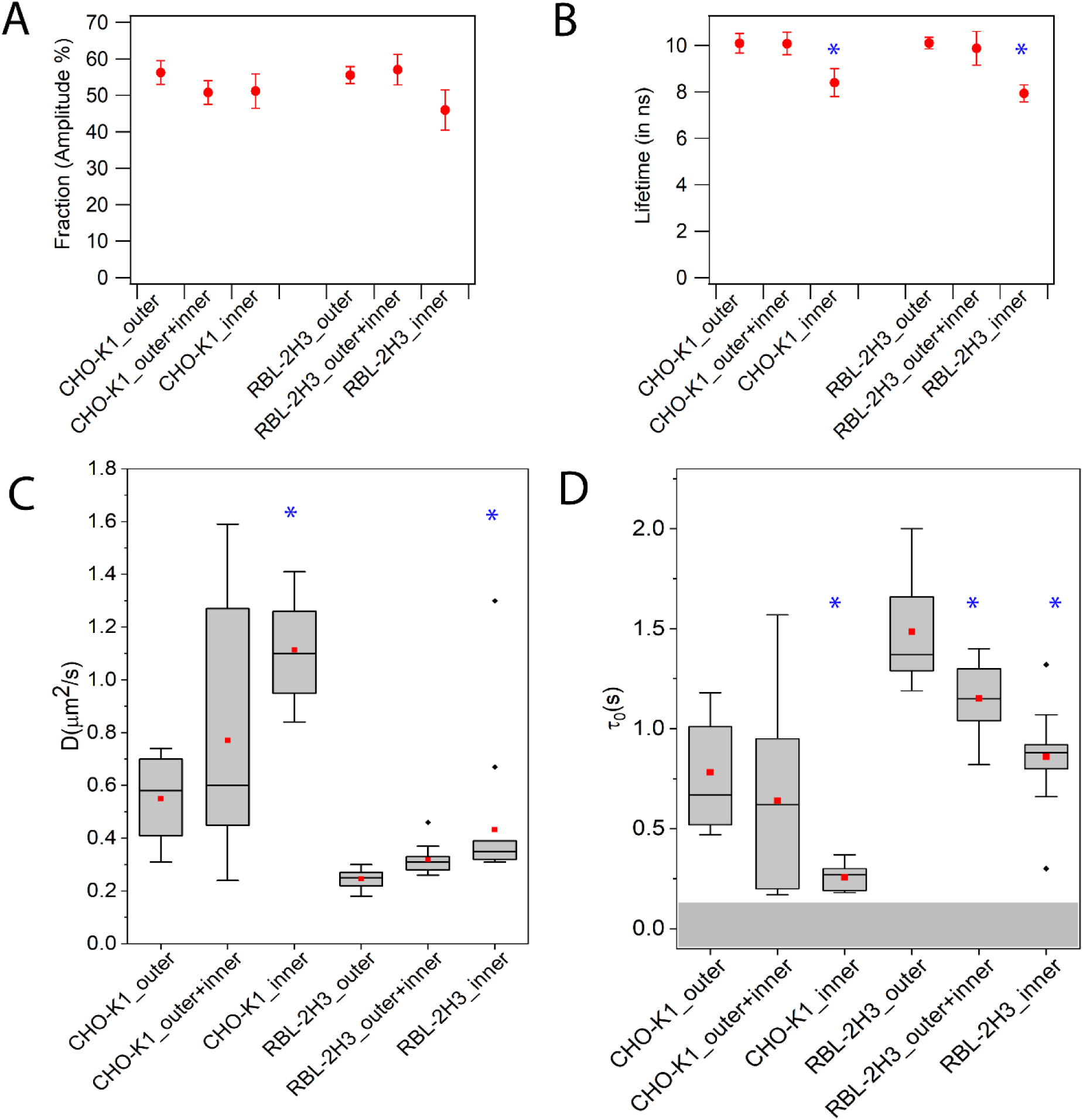
Leaflet specific FLIM and ITIR-FCS analysis of fluorescently labeled phosphatidylcholine analogs in CHO-K1 and RBL-2H3 cells at room temperature. (A) Fraction percentage (amplitude) of NBD-PC long lifetime. (B) Long lifetime of NBD-PC measured in nanoseconds. Since out of the three lifetimes obtained for each measurement only the two longer lifetimes are sensitive to the membrane packing, for clarity we show only the long component. (C) Average diffusion coefficient of TopFluor-PC. (D) FCS diffusion law intercept of TopFluor-PC. Data is pooled average, and error bars are standard deviation. ITIR-FCS data is represented as means of boxplots. Maximum and minimum are shown as vertical boxes with error bars, the 1st, and 99th percentiles are shown as (x). Means are shown as red squares. To ensure reproducibility, FLIM experiments have been repeated at least three times independently. ITIR-FCS data is the average of at least 12 measurements each done on different cells from three independent experiments. (*) indicates the significant differences with respect to the values measured in outer leaflet (paired t-test, p-value < 0.05). For representative raw data refer supplementary information figure S2.

The higher domain fraction in RBL-2H3 cells compared to CHO-K1 cells has been addressed in our previous studies^36,37^. Lower lipid mobility and more significant entrapment of the probe can be attributed to higher levels of sphingolipids in the RBL-2H3 plasma membrane which is known to form trapping sites in the cell membrane^39^. This is also supported by an overall longer lifetime of NBD lipid analogues in giant plasma membrane vesicles (GPMVs) derived from RBL-2H3 cells than CHO-K1 cells (Figure S3) measured at 25 °C and 37 °C. Since GPMVs preserve membrane composition but not organization^40^, the long lifetime in RBL-2H3 GPMVs show that the RBL-2H3 cell membrane composition confers a more ordered environment and a higher domain fraction.

### Leaflet specific analysis of sphingomyelin fluorescent analogues in CHO-K1 and RBL-2H3 cell membranes

The existence of cholesterol-sphingomyelin complexes in cell membranes has been demonstrated previously^41–43^. It is expected that the organization and dynamics of the molecules forming these nanoscale assemblies would differ from that of PC which is ubiquitously present in the cell membrane. To understand the asymmetric transbilayer organization of domain-specific molecules, we characterized the environment and dynamics of SM fluorescent lipid analogues. Sphingomyelins are the most abundant sphingolipids present in the plasma membrane of mammalian cells^11^.

They consist of a ceramide backbone and phosphocholine headgroup exhibiting a narrower cylindrical geometry than phosphatidylcholine and a phase transition above room temperature and thus, exist in the gel phase at room temperature. Sphingomyelins interact with cholesterol to form lipid domains in the membranes, but can also exist freely^41,44,45^. They are predominantly present in the outer leaflet; however, there is some evidence suggesting a pool of SM localized in the inner leaflet^46,47^. Overall, this is consistent with our quenching experiments that show ∼ 80% SM lipid analogues are present in the outer leaflet (Table S1).

The use of tail-labeled NBD lipid analogues for membrane studies is debatable as they may not behave like the endogenous lipids^48,49^ Therefore, to validate that the NBD probes are sensitive to cholesterol content we performed methyl-β-cyclodextrin induced cholesterol depletion experiments. We observed that as expected, the effect of cholesterol depletion significantly affects the NBD-SM lifetime as indicated by a drop of 6 ns in both cell lines (Figure S4). However, the NBD-PC lifetime shows a decline of 1 ns only. This experiment verifies that the disruption of cholesterol domains directly influences the SM microenvironment but to a much lesser extent that of PC. Thus, NBD lipid analogues reside in environments similar to their endogenous forms and therefore, can be used to examine the domain-specific plasma membrane organization.

Next, we proceeded with the leaflet specific analysis of SM probes. The lifetime analysis of NBD-SM immediately after labeling i.e. in the outer leaflet shows a lifetime of around 10.5 ns in both cell lines (Figure 3 *A, B*) suggesting the existence of ordered domains surrounding the NBD-SM molecules in the outer leaflet. ITIR-FCS results show that in case of TF-SM, *D* in the outer leaflet is around two-fold higher in CHO-K1 cells (0.45 ± 0.07 μm^2^/s) than in RBL-2H3 (0.22 ± 0.03 μm^2^/s) (Figure 3 *C, D*). As expected, the *τ0* is positive in the outer leaflet of both cells with a higher value in RBL-2H3 cells (*τ0* = 1.25 ± 0.25 s) than in CHO-K1 (*τ0* = 0.59 ± 0.12 s) consistent with the existence of a fraction of lo phase in the outer leaflet. After transbilayer redistribution of the probe between both leaflets (Fig. 1B, middle), the lifetime of NBD-SM was 0.5 ns shorter than in the outer leaflet of CHO-K1 cells. However, in RBL-2H3 cells there was a 1 ns increase in the lifetime of the probe implying that post redistribution NBD-SM localizes in more ordered microenvironments. ITIR-FCS measurements show for TF-SM a 40 % increase of *D* and a drop of *τ0* in CHO-K1 cells suggesting faster lipid mobility in the inner leaflet while in RBL-2H3 cells, there is no significant difference in the lipid mobility and probe entrapment. Subsequently, the analysis of NBD-SM exclusively on the inner leaflet confirmed the existence of an ld phase in the inner leaflet of CHO-K1 cells as shown by a fluorescence lifetime of 7.80 ± 0.55 ns and an even more ordered inner leaflet in RBL-2H3 cells (lifetime of 9.1 ± 0.04 ns). The fraction of the long lifetime in the inner leaflet is reduced to ∼30% with respect to 50% of the outer leaflet for both cell lines. Consistent with lifetime measurements, TF-SM shows in the inner leaflet a 36 % higher *D* relative to that in the outer leaflet and *τ0* was reduced from 0.59 ± 0.12 s to 0.38 ± 0.13 s in the inner leaflet of CHO-K1 cells. On the contrary in RBL-2H3 cells there is no significant difference in *D* when comparing inner and outer leaflet. Moreover, the change in *τ0* shows no significant effect (1.25 ± 0.25 s to 1.08 ± 0.23 s) as indicated by the t-test (p-value < 0.05). In contrast to PC, SM lipid analogues in RBL-2H3 cells indicate a higher domain fraction in the inner leaflet of the plasma membrane leading to slow diffusion and longer lifetime.

**Figure 3:**
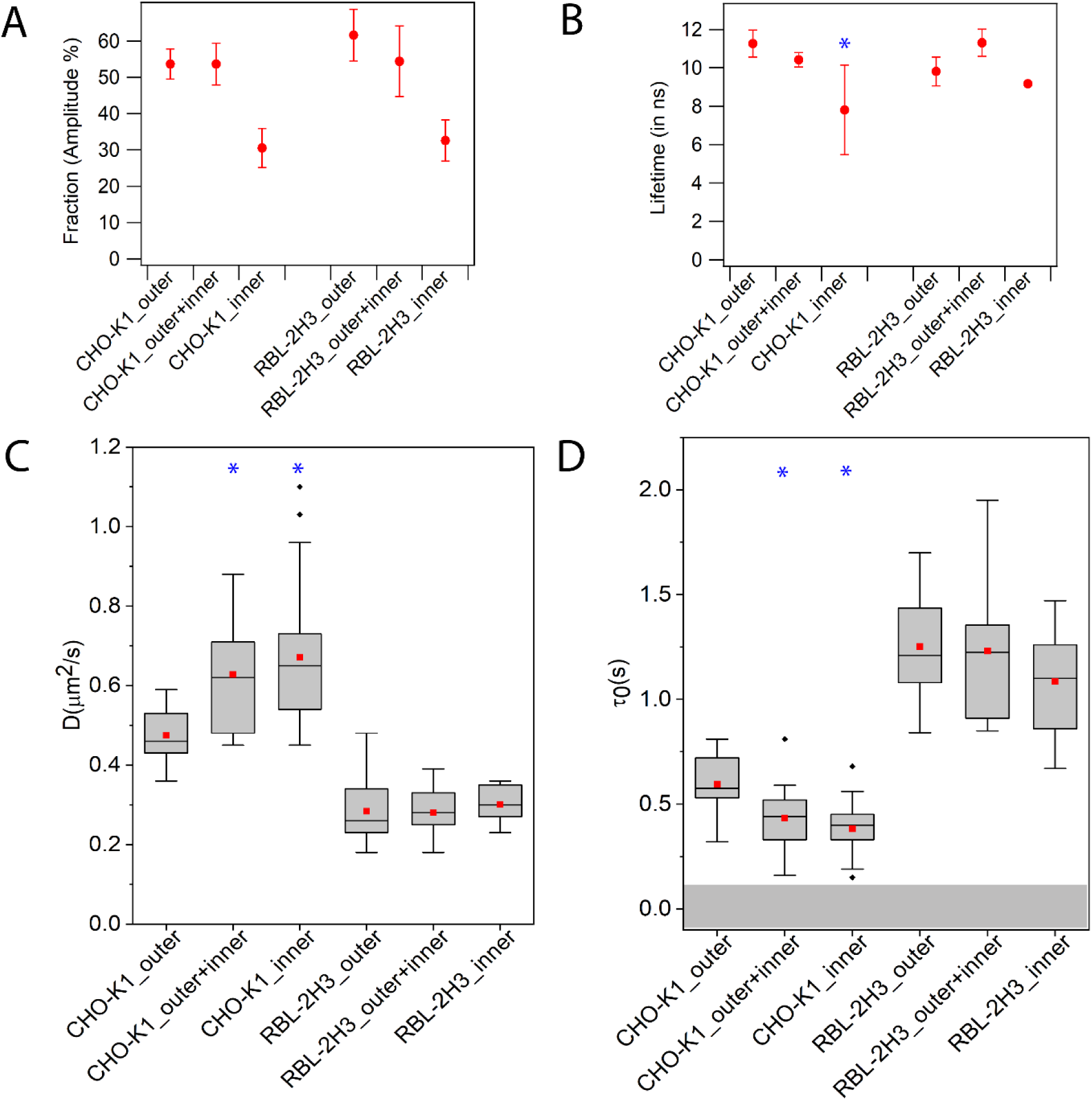
Leaflet specific FLIM and ITIR-FCS analysis of fluorescently labeled sphingomyelin analogs in CHO-K1 and RBL-2H3 cells at room temperature. (A) Fraction percentage (amplitude) of NBD-SM long lifetime. (B) Long lifetime of NBD-SM measured in nanosecond. Only long lifetime shown (See legend to figure 2). (C) Average diffusion coefficient of TopFluor-SM. (D) FCS diffusion law intercept of TopFluor-SM. Data is pooled average, and error bars are standard deviation. ITIR-FCS data is represented as means of boxplots. Maximum and minimum are shown as vertical boxes with error bars, the 1st, and 99th percentiles are shown as (x). Means are shown as red squares. To ensure reproducibility, FLIM experiments have been repeated at least three times independently. ITIR-FCS data is the average of at least 12 measurements each done on different cells from three independent experiments. (*) indicates the significant differences with respect to the values measured in outer leaflet (paired t-test, p-value < 0.05). For representative raw data refer supplementary information figure S5.

Our results reveal differences in individual leaflet packing in the two cell lines. CHO-K1 cells show a more profound difference between the outer and inner leaflet environment than RBL-2H3 cells. This could imply the occurrence of higher dynamic coupling between the two leaflets in RBL-2H3 cells. Unlike PC analogues, SM analogues manifest similar dynamics and organization when compared between the two leaflets. Previous studies suggest that interdigitation of SM can mediate interleaflet coupling in the plasma membrane^50^. Similar dynamics of SM probes in the two leaflets could support the proposition of SM mediated inter leaflet coupling. In addition to that, our readouts obtained on SM probes also show that RBL-2H3 cells manifest a more ordered membrane environment than CHO-K1 cells, as already noted in the case of PC analogues.

In summary, besides demonstrating a lower degree of heterogeneity in the inner leaflet, our results also demonstrate that the asymmetric arrangement of the plasma membrane can vary for different cell types used. Furthermore, probe related differences suggest that different aspects of organisation and dynamics are recorded. This can be visualized by analysing the differences in the ratios of *D* measured in the inner leaflet and the outer leaflet which also demonstrate the probe and cell type specific plasma membrane asymmetry (Table S2).

### Leaflet specific analysis of phosphatidylserine fluorescent analogues in CHO-K1 and RBL-2H3 cell membranes

We probed the organization and dynamics of fluorescent phosphatidylserine analogues, to characterize the inner leaflet organization^51^. Usually, it comprises the fluid fraction of the plasma membrane^52^ however, there is evidence suggesting its direct interaction with the cytoskeleton due to which PS can exhibit lower mobility^53^. Close proximity of the inner leaflet with the cytoskeletal network could also alter the mode of lipid diffusion as molecules hindered by meshworks are shown to undergo hop diffusion^54^ and additional protein interactions in the inner leaflet could result in distinct lipid microenvironments^55^.

First, we estimated the percentage of PS probes residing in the inner leaflet by performing outer leaflet fluorescence quenching experiments. In CHO-K1 cells, around 70% PS fluorescent analogues reside in the inner leaflet while in RBL-2H3 cells, 80% of probe molecules are in the inner leaflet (Table *S1*).

When measured in the outer leaflet, NBD-PS shows a lifetime of 10.21 ± 0.62 ns in CHO-K1 cells while 8.98 ± 0.29 ns in RBL-2H3 cells (Figure 4 *A, B*) indicating the existence of an lo phase in the outer leaflet. Unlike PC and SM analogues where the occurrence of an lo phase correlates with slower lipid mobility, TF-PS diffuses two-fold faster and shows lower transient entrapment than the TF-PC and TF-SM in the respective cell lines measured under the same experimental conditions (Figure 4 *C, D*). In this case also, *D* of TF-PS is around 2.5 times higher in CHO-K1 cells (1.1 ± 0.26 μm^2^/s) than in RBL-2H3 cells (0.47 ± 0.13 μm^2^/s).

**Figure 4:**
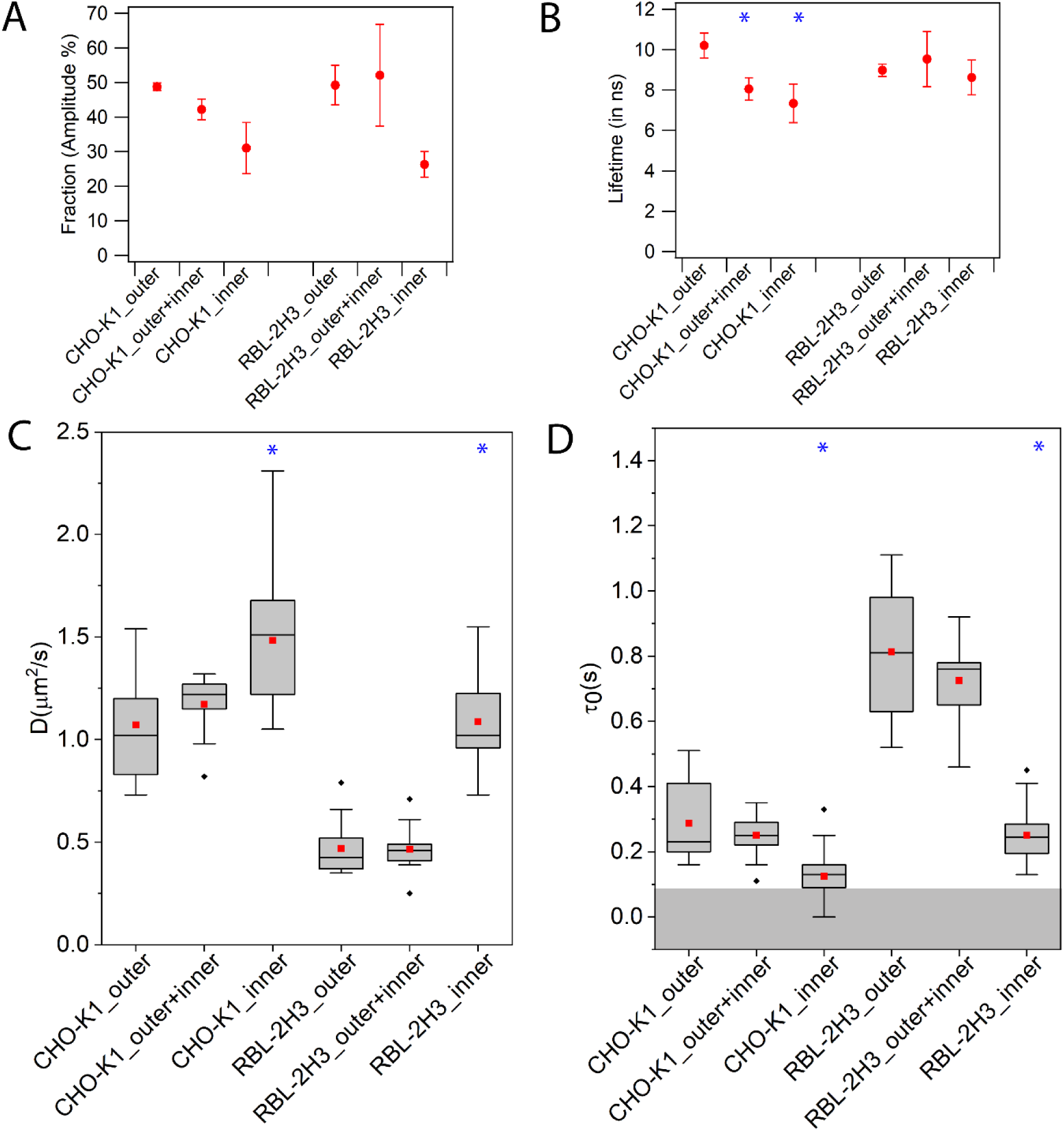
Leaflet specific FLIM and ITIR-FCS analysis of fluorescently labeled phosphatidylserine analogs in CHO-K1 and RBL-2H3 cells at room temperature. (A) Fraction percentage (amplitude) of NBD-PS long lifetime. (B) Long lifetime of NBD-PS measured in nanosecond. Only long lifetime shown (See legend to figure 2). (C) Average diffusion coefficient of TopFluor-PS. (D) FCS diffusion law intercept of TopFluor-PS. Data is pooled average, and error bars are standard deviation. To ensure reproducibility, FLIM experiments have been repeated at least three times independently. ITIR-FCS data is represented as means of boxplots. Maximum and minimum are shown as vertical boxes with error bars, the 1st, and 99th percentiles are shown as (x). Means are shown as red squares. ITIR-FCS data is the average of at least 12 measurements each done on different cells from three independent experiments. (*) indicates the significant differences with respect to the values measured in outer leaflet (paired t-test, p-value < 0.05). For representative raw data refer supplementary information figure S6.

Moreover, transient entrapment of the probe in domains is significantly lower in CHO-K1 cells (*τ*_*0*_ = 0.27 ± 0.12 s) than in RBL-2H3 cells (*τ*_*0*_ = 0.81 ± 0.19 s). A combined readout from the two leaflets shows a 2 ns drop of the NBD-PS lifetime in CHO-K1 cells, but no significant change is observed in RBL-2H3 cells. However, the TF-PS dynamics remain largely unaltered for both cell lines when the two leaflets are analyzed together. Finally, on probing the inner leaflet exclusively, we observed a 2.3 ns lower lifetime of NBD-PS in CHO-K1 inner leaflet relative to the outer leaflet while it remains the same in RBL-2H3 cells. Moreover, the fraction of the long lifetime component originating from NBD-PS is reduced to 30% in both cell lines. Consistent with this trend, a lowering of the NBD-PS lifetime was accompanied with a 26% faster diffusion of TF-PS and *τ*_*0*_ close to the threshold for free diffusion in CHO-K1 cells. Despite the same lifetime in RBL-2H3 cells, we observed a 120% higher *D* (0.47 ± 0.13 μm^2^/s to 1.08 ± 0.23 μm^2^/s) in the inner leaflet of the membrane and *τ*_*0*_ decline from 0.73 ± 0.19 s in the outer leaflet to 0.22 ± 0.08 s in the inner leaflet. This could mean that the differences in the microenvironment of NBD-PS in the two leaflets of RBL-2H3 cell membranes is below the detection limits of FLIM. In Hela cells also, NBD-PS and NBD-PC lifetime was measured to be similar^16^. In this regard, it is important to consider that in addition to membrane packing, NBD lifetime is sensitive to environmental polarity. Moreover, NBD probes have been shown to exhibit red edge excitation shift (REES) when NBD lipid probes loop back at the water-membrane interface^25,56^. Due to the multiple factors that influence the NBD lifetime, evaluation of NBD lifetime alone can result in misleading interpretations, and it is essential to supplement the lifetime results with another quantitative method such as ITIR-FCS as used in this study.

These observations indicate an overall higher molecular mobility of PS lipid analogues in cell membranes compared to other choline lipids. In CHO-K1 cells, PS lipid analogues show a distinct microenvironment in the two leaflets as assessed by the NBD lifetime and TF-PS diffusion. Moreover, while in the inner leaflet PS lipid analogues reside in a more fluid environment as revealed by lifetime characteristic of ld phase vesicles (DOPC), higher mobility and free diffusion of the probe. RBL-2H3 cells, on the other hand, do not show differences in NBD-PS environment in the two leaflets. However, significant differences in the diffusion properties of TF-PS in the two leaflets suggest a less viscous environment in the inner leaflet^57^. Despite the lower membrane viscosity, factors such as cytoskeleton network and protein interactions could alter the solvent polarity and tendency of NBD probes to loop back which can contribute to the long NBD lifetimes in the inner leaflet.

## CONCLUSIONS

The plasma membrane is structurally complex and dynamic with a diversity of lateral molecular movements ranging from directed diffusion of immobile protein-lipid clusters to free diffusion of molecules and transversal movements of lipids occurring at varying rates.

Besides, there are fluctuating nanoscale assemblies on the membrane scaling from 5 nm to 500 nm. To understand the organization and dynamics of such a system, it is essential to use methods with varying spatial and temporal resolutions. In this study, we performed a spatiotemporal analysis of the plasma membrane asymmetry in live mammalian cells using advanced fluorescence methods with multiplexing capabilities, namely, FLIM and ITIR-FCS. These methodologies enable us to distinguish domain/leaflet specific organization and dynamics even across closely related cell lines. We characterize the plasma membrane asymmetry by evaluating the microenvironment and dynamics of fluorescent analogues of PC, SM, and PS which exhibit specific preferences for domains and leaflets.

By using a combinatorial approach of FLIM and ITIR-FCS we can sense the microenvironment independent of concentration and mobility (FLIM) while at the same time measuring membrane dynamics and organization (ITIR-FCS). Our results show that in general in mammalian cell membranes, the outer leaflet is more densely packed than the inner leaflet as demonstrated by a longer lifetime, slower probe mobility and greater transient confinement of lipids (Figure 2,3,4) in the former than the latter. However, the detailed membrane dynamics and organization reveals cell line specific differences in the plasma membrane asymmetry. For instance, in RBL-2H3 cells, SM analogues showed similar membrane packing and diffusion properties in both leaflets (Figure 3 C, D) while in CHO-K1 cells clear differences for SM analogues were detected between both leaflets possibly implying strong interleaflet coupling in RBL cells. Furthermore, accessible details of membrane dynamics and organization depend also on the probes used as shown by probe specific readouts. Overall our study demonstrates how FLIM and ITIR-FCS can analyse differences in dynamics and organization between outer and inner plasma membrane leaflets as seen by different probe molecules in dependence of the cell line investigated.

## ACKNOWLEDGMENTS

AG is the recipient of the research scholarship from the National University of Singapore (NUS). TW and AH acknowledge funding by the HU-NUS joint cooperation grant 2017.

## Abbreviations

The abbreviations used are:

ITIR-FCS: Imaging total internal spectroscopy fluorescence correlation spectroscopy;
ACF: autocorrelation function;
FLIM: Fluorescence lifetime imaging microscopy;
FLS: fluorescence lifetime spectroscopy;
GUV: giant unilamellar vesicle;
GPMV: giant plasma membrane vesicle;
NBD: 7-nitro-2-1,3-benzoxadiazol-4-yl)amino]hexanoyl;
TF: TopFluor;
PC: phosphatidylcholine;
SM: sphingomyelin;
PS: phosphatidylserine;
EMCCD: Electron-multiplying charge coupled device;
TCSPC: Time-correlated single-photon counting.

## FOOTNOTES

### AUTHOR CONTRIBUTIONS

A.H. and T.W supervised the project. A.G., T.K. performed the FLIM experiments. A.G. performed the ITIR-FCS experiments. A.G. performed the data analysis. All authors contributed to the manuscript writing.

