## Supplementary file 1 for "Analysing plasma membrane asymmetry of lipid organisation by fluorescence lifetime and correlation spectroscopy"

### SUPPLEMENTARY INFORMATION

#### *S1: Validation of NBD lifetime sensitivity for membrane phase behaviour studies*

To estimate the range of NBD lifetimes expected in the membrane environment we measured the lifetime of NBD-PC in vesicles composed of well-defined lipid compositions and known phases. To demonstrate the sensitivity of NBD lifetime to membrane packing, the measurements were done at a temperature range of 25 °C - 45 °C as temperature alters the membrane packing.

The long lifetime of NBD-PC in DOPC LUV shows a decrease in the lifetime (6.81ns -5.8 ns) with an increase in temperature in the mentioned range (Figure S1). A similar decreasing trend was observed for DOPC:SSM:Chol (2:2:6) LUVs with a long lifetime value being higher for this lipid composition than pure DOPC LUVs. The lifetime drops from 9.81 ns to 7.07 ns at a temperature range of 25 °C - 45 °C. In the case of DOPC:SSM:Chol (1:1:1) LUVs, the two long lifetimes show a decrease in the lifetime with increasing temperature, the longer one decreasing from 9.61 ns to 7.87 ns and the shorter one varies from 5.06 ns to 4.21 ns. The lowering of NBD lifetime upon temperature increase is expected as at higher temperatures, the overall membrane packing becomes loose and the increase in the non-radiative decay of NBD[25]. The longer lifetime of NBD-PC observed in the case of DOPC:SSM:Chol (2:2:6) with respect to that is obtained in pure DOPC is attributed to the more ordered lipid packing in the former than the latter. These results provide the range of NBD lifetime variation in membranes with extreme phase behaviours. This information will serve as the reference while interpreting the cell membrane experiments.

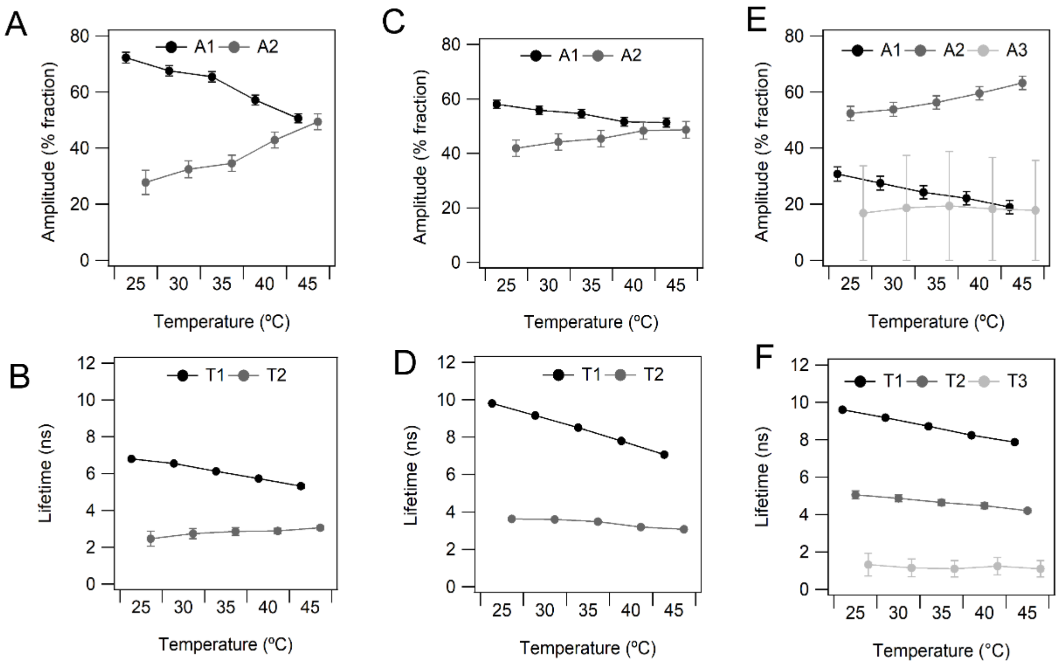


**Figure S1:** Fractional amplitudes and Fluorescence lifetimes τ of $C_6$-NBD-PC in large unilamellar vesicles at various temperatures measured using fluorescence lifetime spectroscopy (A,B) Pure DOPC exhibiting liquid disordered phase (C,D) DOPC:SSM:Chol (2:2:6) exhibiting liquid ordered phase (E,F) DOPC:SSM:Chol (1:1:1) exhibiting coexisting liquid ordered and disordered phase. (refer the mentioned articles for the phase specific lipid compositions)[34,35].

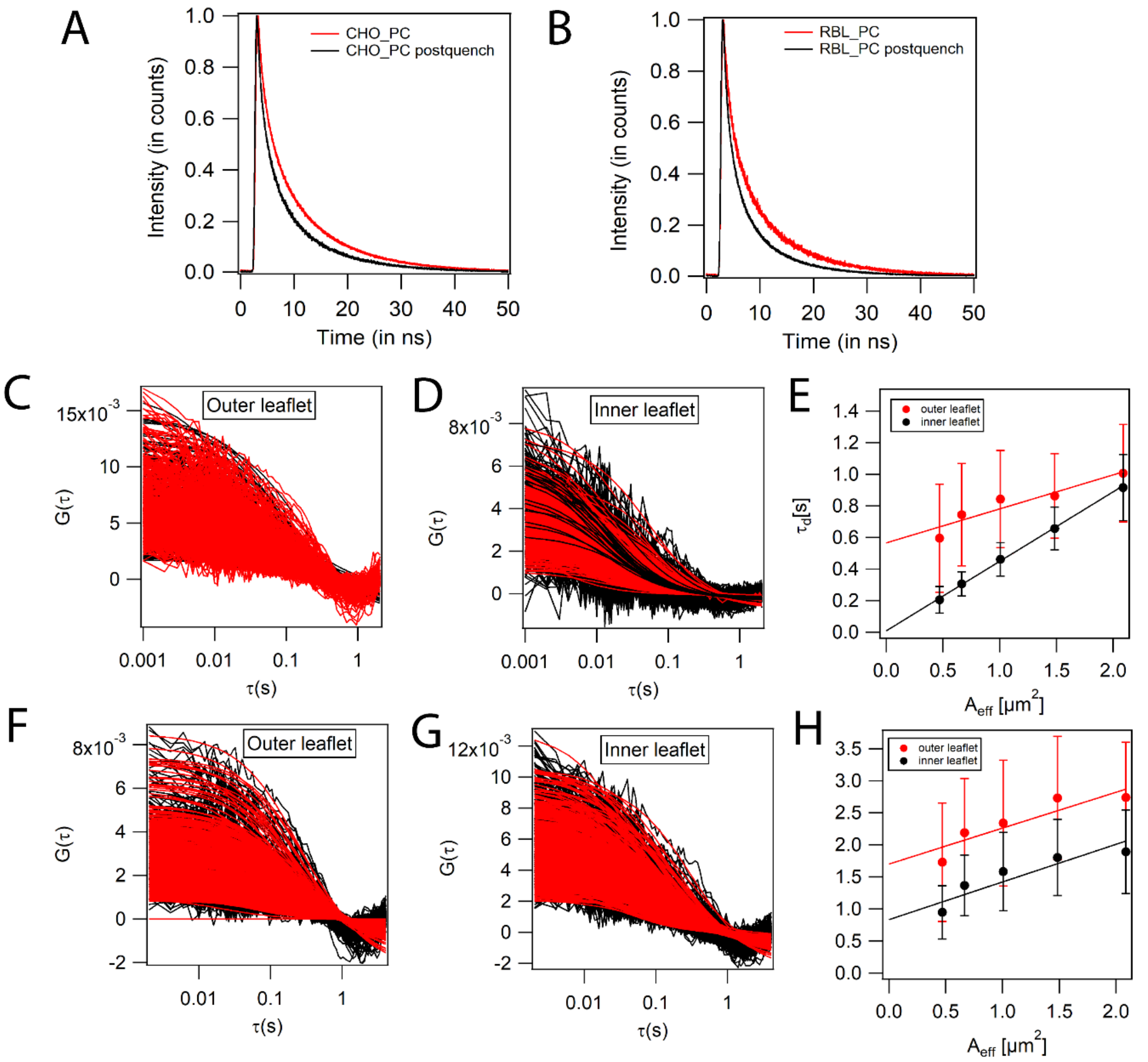


**Figure S2:** Representative raw data for FLIM and ITIR-FCS experiments to analyze leaflet specific dynamics and organization of phosphatidylcholine fluorescent analogues A) Averaged intensity decay of NBD-PC in the outer and inner leaflet of CHO-K1 cell membrane. B) Averaged intensity decay of NBD-PC in the outer and inner leaflet of RBL-2H3 cell membrane. C) Autocorrelation curves for TopFluor-PC labeling the outer leaflet of CHO-K1 cell membrane D) Autocorrelation curves for TopFluor-PC labelllabeling the inner leaflet of CHO-K1 cell membrane (E) FCS diffusion law plots for TopFluor-PC diffusion in outer and the inner leaflet of CHO-K1 cell membrane. F) Autocorrelation curves for TopFluor-PC labeling the outer leaflet of RBL-2H3 cell membrane G) Autocorrelation curves for TopFluor-PC labeling the inner leaflet of RBL-2H3 cell membrane (H) FCS diffusion law plots for TopFluor-PC diffusion in outer and the inner leaflet of RBL-2H3 cell membrane.

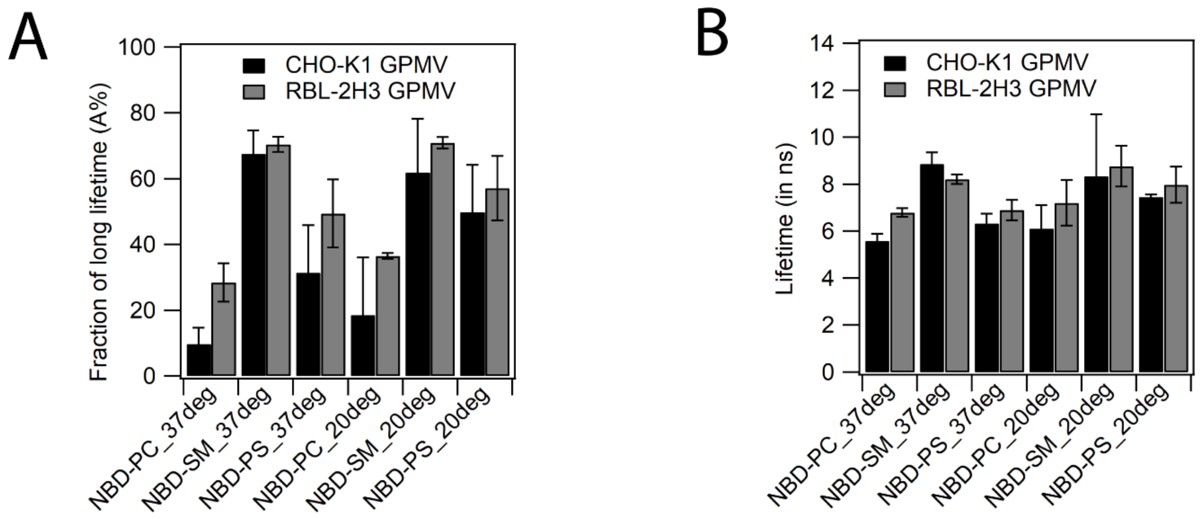


**Figure S3:** Comparison of fluorescence lifetimes of $C_6$-NBD analogues (PC, SM and PS) in GPMVs derived from CHO-K1 and RBL-2H3 cells. The measurements are performed at 20 °C and 37 °C. Error bars represent the standard deviation. (n = 3)

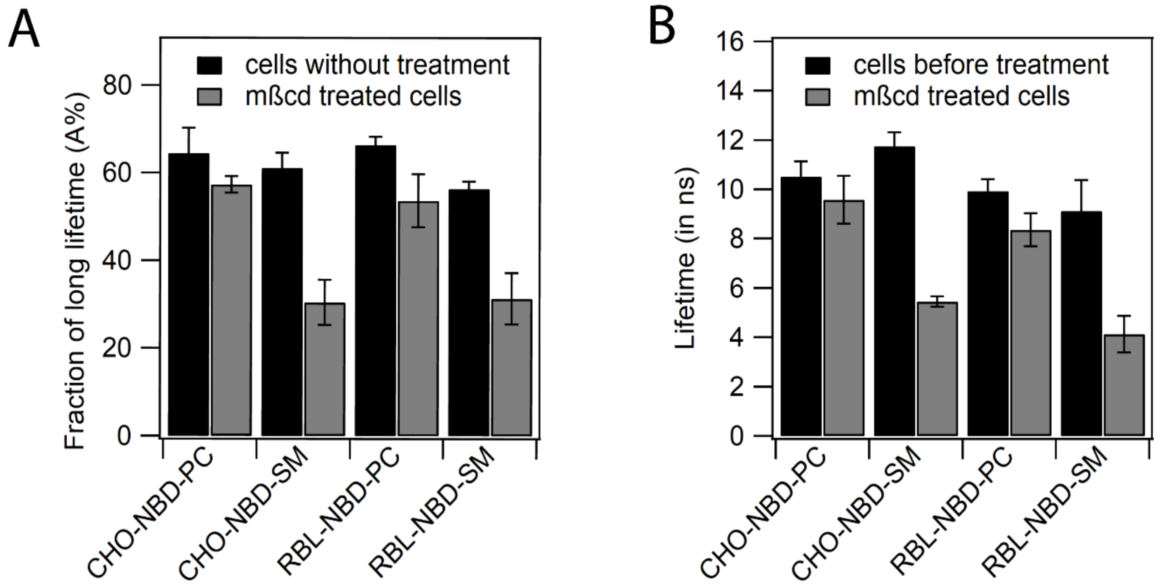


**Figure S4:** Validation of NBD lipid analogue (NBD-PC and NBD-SM) localization in CHO-K1 and RBL-2H3 cell membranes using methyl-β-cyclodextrin depletion experiments. (n = 3) (A) Fractional amplitude of long component (B) Fluorescence lifetime of the long lifetime. Error bars represent the standard deviation.

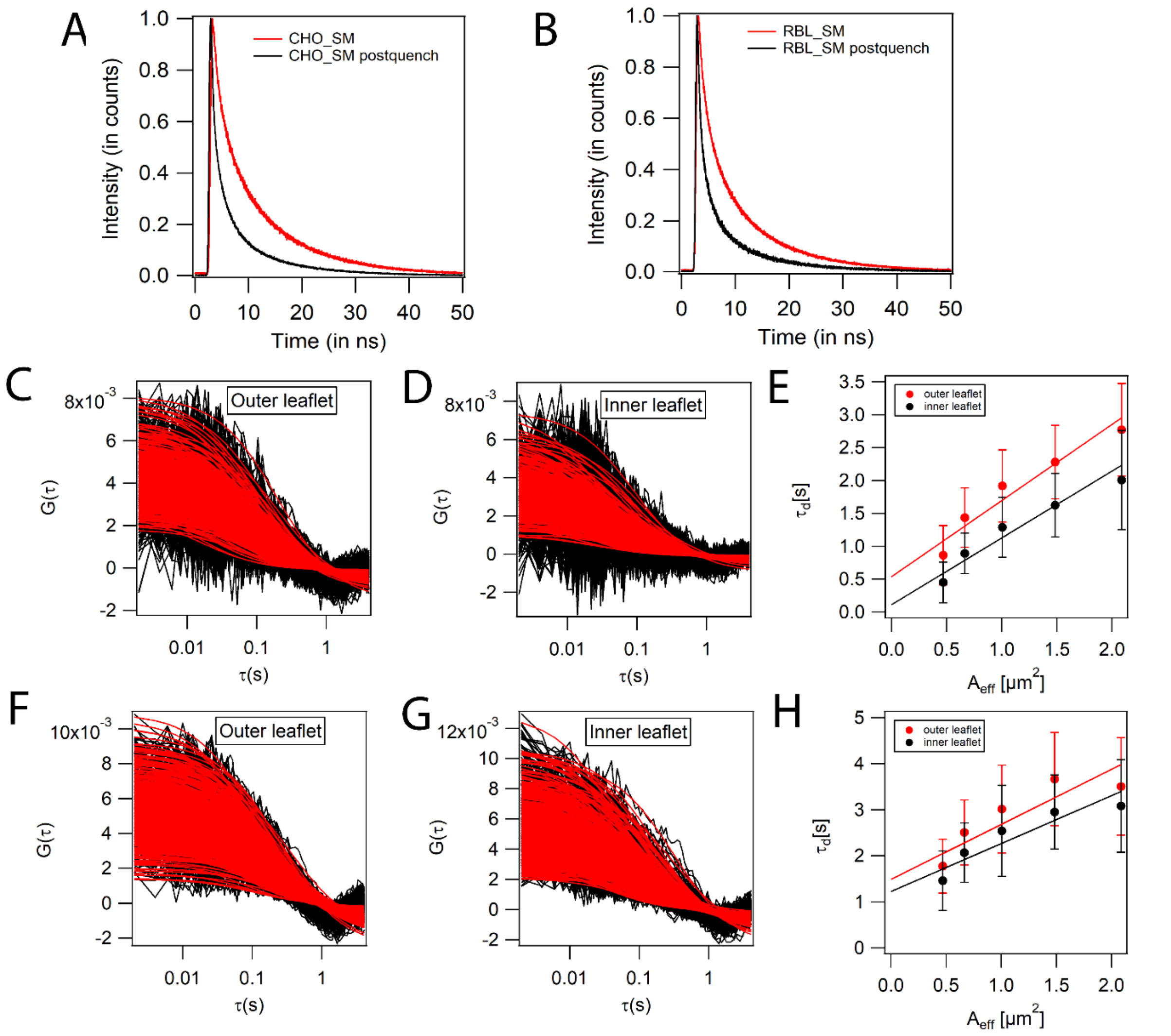


**Figure S5:** Representative raw data for FLIM and ITIR-FCS experiments to analyze leaflet specific dynamics and organization of sphingomyelin fluorescent analogues A) Averaged intensity decay of NBD-SM in the outer and inner leaflet of CHO-K1 cell membrane. B) Averaged intensity decay of NBD-SM in the outer and inner leaflet of RBL-2H3 cell membrane. C) Autocorrelation curves for TopFluor-SM labeling the outer leaflet of CHO-K1 cell membrane D) Autocorrelation curves for TopFluor-SM labeling the inner leaflet of CHO-K1 cell membrane (E) FCS diffusion law plots for TopFluor-SM diffusion in outer and the inner leaflet of CHO-K1 cell membrane. F) Autocorrelation curves for TopFluor-SM labeling the outer leaflet of RBL-2H3 cell membrane G) Autocorrelation curves for TopFluor-SM labeling the inner leaflet of RBL-2H3 cell membrane (H) FCS diffusion law plots for TopFluor-SM diffusion in outer and the inner leaflet of RBL-2H3 cell membrane.

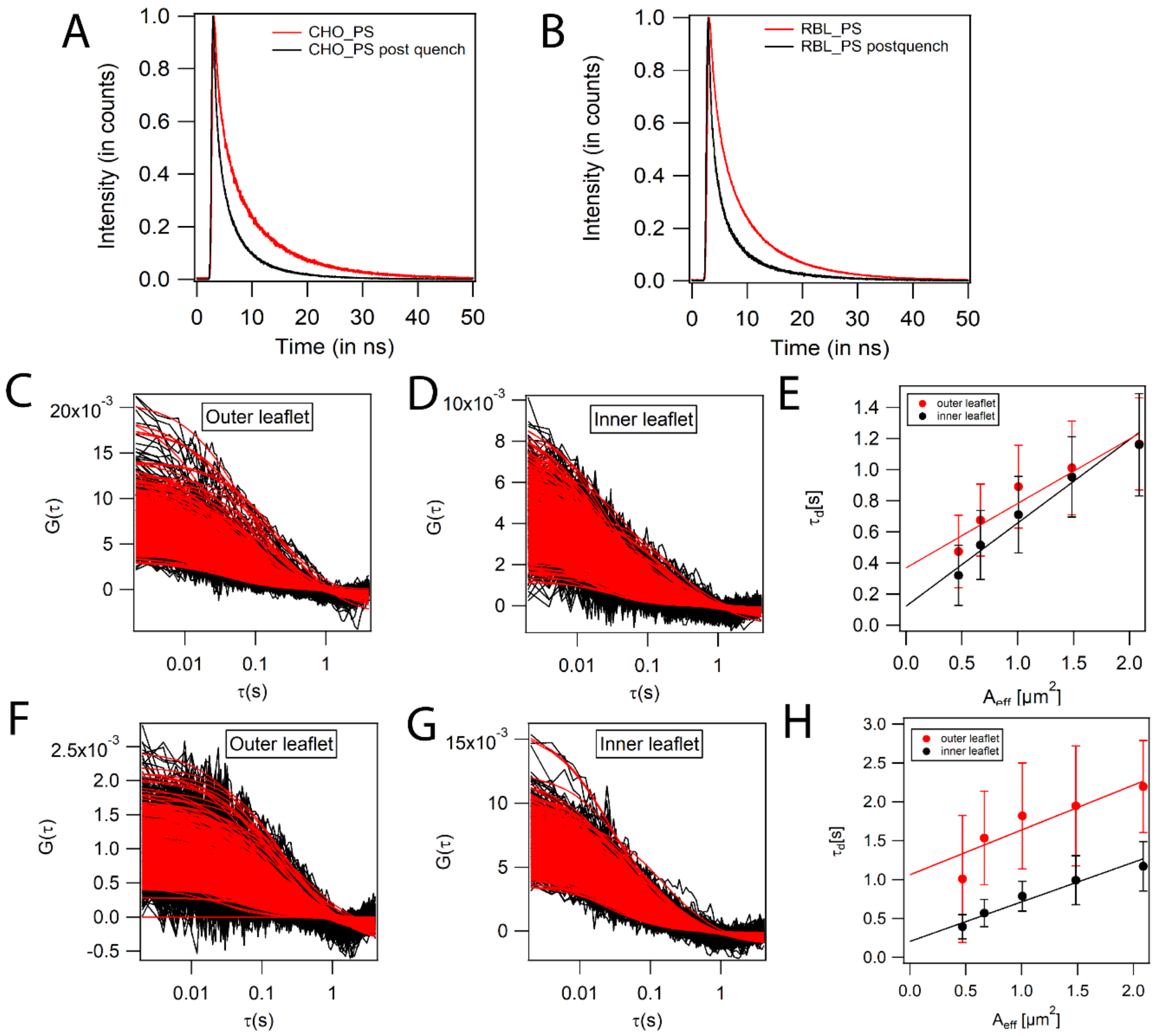


**Figure S6:** Representative raw data for FLIM and ITIR-FCS experiments to analyze leaflet specific dynamics and organization of phosphatidylserine fluorescent analogues A) Averaged intensity decay of NBD-PS in the outer and inner leaflet of CHO-K1 cell membrane. B) Averaged intensity decay of NBD-PS in the outer and inner leaflet of RBL-2H3 cell membrane. C) Autocorrelation curves for TopFluor-PS labeling the outer leaflet of CHO-K1 cell membrane D) Autocorrelation curves for TopFluor-PS labeling the inner leaflet of CHO-K1 cell membrane (E) FCS diffusion law plots for TopFluor-PS diffusion in outer and the inner leaflet of CHO-K1 cell membrane. F) Autocorrelation curves for TopFluor-PS labeling the outer leaflet of RBL-2H3 cell membrane G) Autocorrelation curves for TopFluor-PS labeling the inner leaflet of RBL-2H3 cell membrane (H) FCS diffusion law plots for TopFluor-PS diffusion in outer and the inner leaflet of RBL-2H3 cell membrane.

| Fluorescent probe | Quenching % (CHO-K1) | Quenching % (RBL-2H3) |
|---|---|---|
| NBD-PC | 66.56 ± 2.35 | 77.12 ± 1.52 |
| NBD-SM | 82.23 ± 2.71 | 84.81 ± 0.91 |
| NBD-PS | 32.12 ± 2.43 | 18.12 ± 4.32 |
| TF-PC | 85.34 ± 1.45 | 82.32 ± 3.11 |
| TF-SM | 87.21 ± 2.43 | 88.12 ± 1.21 |
| TF-PS | 29.16 ± 4.21 | 16.43 ± 3.82 |

**Table S1:** Quantification of the amount of fluorescent lipid analogues accessible towards quenching agent after the incubation of labeled cells for 2 hours. The fluorescence intensity was measured 2 hours after the labeling before and after the quenching. NBD lipid analogues were quenched by 25 mM sodium dithionite and TopFluor lipid analogues are quenched by 50 μM 16-Doxyl lipids. Means and standard deviations are given (n = 3).

| | $D_{inner}/D_{outer}$ TF-PC | $D_{inner}/D_{outer}$ TF-SM | $D_{inner}/D_{outer}$ TF-PS |
|---|---|---|---|
| CHO-K1 | 2.02 | 1.41 | 1.36 |
| RBL-2H3 | 1.76 | 1.05 | 2.31 |

**Table S2 :** Ratio of inner and outer leaflet diffusion coefficients of TF-(PC, SM and PS) in CHO-K1 and RBL-2H3 cells.
